# Mouse dehydration as a consequence of water bottle design

**DOI:** 10.64898/2026.06.15.732413

**Authors:** Fritz Kutschera, Sebastian Lindner, Thomas G. Oertner

## Abstract

Animal care technicians reported that accidental deaths of mice occur more frequently under certain weather conditions. This is surprising, given that modern mouse facilities are fully climate-controlled. Our investigation revealed that air bubbles become trapped in the drinking tube of water bottles and block the downward flow of water. Mice cannot drink from such bottles and become dehydrated. A coiled wire inside the drinking tube dislodges the air bubbles and ensures unrestricted access to water.

## Main

Research institutions invest considerable resources in animal facilities to ensure the wellbeing of their animals. Healthy experimental animals are essential for high quality, reproducible science, and mice account for 73% of the laboratory animals used in Germany^1^. Despite their best efforts, however, accidental deaths with no discernible cause do occur in every animal facility. Mice have a very high metabolic rate and cannot tolerate dehydration for very long^2^. To reduce the risk of dehydration, multiple water bottles per cage or additional HydroGel pouches are sometimes used, which should not be necessary in an air-conditioned facility with *ad libitum* water supply. In their guidelines for drinking water supply, the Society of Laboratory Animal Science (GV-SOLAS) states dryly: “However, it can happen that animals die of thirst in front of a full water bottle.”^3^

To study the physics of drinking, we modified a standard drinking bottle by mounting a pressure sensor in its metal cap. We placed the bottle in its normal working position (45°) in a mouse cage. Proximity of the mouse’s snout to the sipper tube was detected by a passive infrared sensor. To detect tongue contact and count licks, we electronically monitored the sipper tube’s resistance to ground (Fig. 1).

**Fig. 1:**
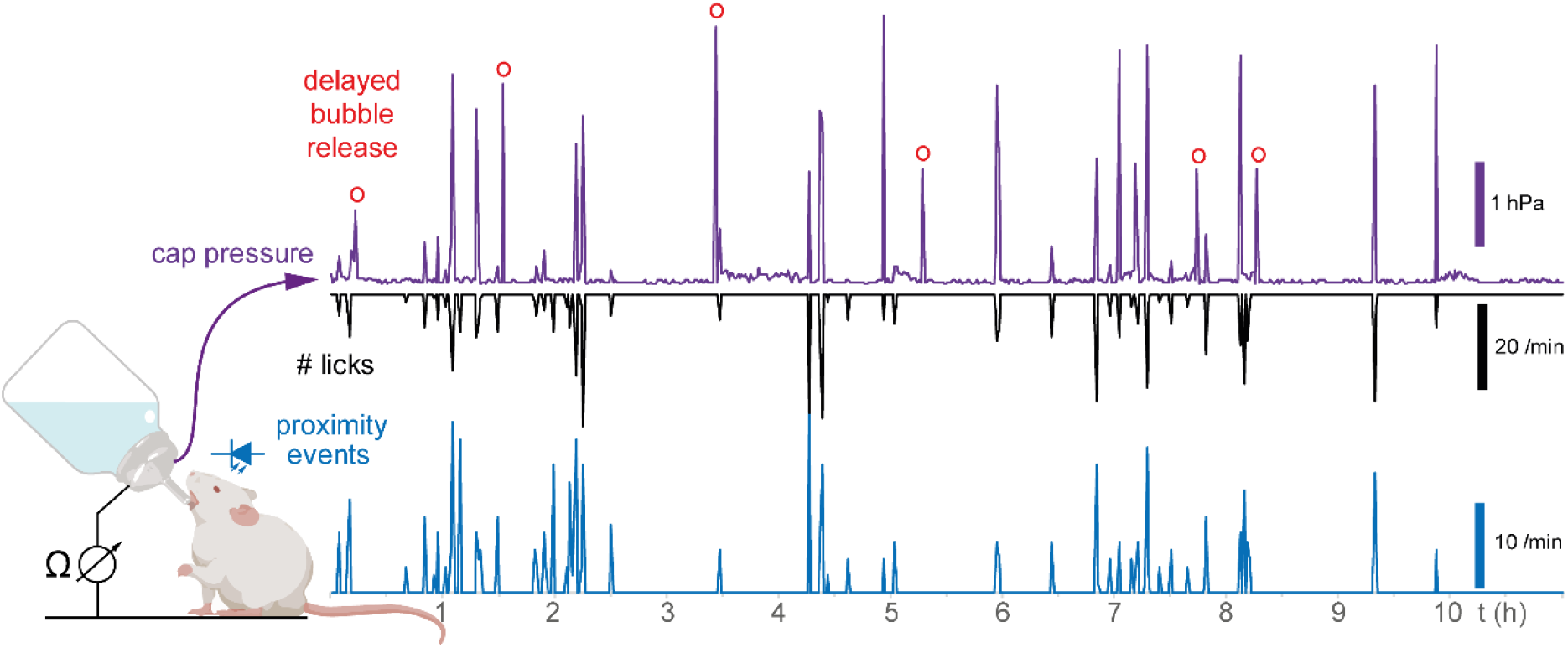
Monitoring the drinking bouts of a single mouse over a 10-hour period. Proximity events (blue trace) coincide with licking periods (black trace). Pressure spikes during licking (violet trace) correspond to air bubbles rising through the water. Pressure spikes without licking (red circles) indicate the spontaneous release of trapped bubbles. The number of events was counted in 1 min windows.

Spikes in bottle cap pressure coincided with tongue contacts, indicating the passage of air bubbles that compensated the pressure difference caused by water extraction. We also noted spontaneous pressure spikes at times when the animal was not in contact with the bottle (Fig. 1, red circles), suggesting that air was transiently trapped inside the cylindrical sipper tube. Mice are not able to suck on the sipper tube; rather, they rapidly cover and uncover the small hole with their tongue, causing water to leak onto their tongue. Extracting water from the bottle creates a partial vacuum in the airspace above the water level. This pressure difference is periodically compensated by air bubbles forming inside the sipper tube and rising inside the bottle. Delayed bubble release events could indicate that pressure equalization is not always instantaneous.

Next, we kept 9 mice separated in Nutrition Monitor™ cages to continuously monitor their activity and water intake over 3 months. We found that mice typically consume the same amount of water every day, distributed over many short visits to the bottle. However, in one occasion, two mice in separate cages simultaneously reduced their water consumption dramatically (Fig. 2). Activity was reduced during this thirst period. On the second day, the mice were motionless for up to 4 hours, indicating severe dehydration, in spite of a well-filled water bottle in each cage. After two days, the mice began to drink again at an elevated rate, starting precisely at the same time. A third mouse showed a similar, but less extreme pattern (Supp. Fig. 1), suggesting that one third of the tested animals was affected. Consulting the weather report, we noted that the start of the thirst period coincided with rapidly increasing atmospheric pressure, and the affected mice became active again and resumed drinking as soon as the pressure dropped. The synchronized behavioral changes suggest that elevated air pressure can somehow compromise the ability of mice to drink from their bottle.

**Fig. 2:**
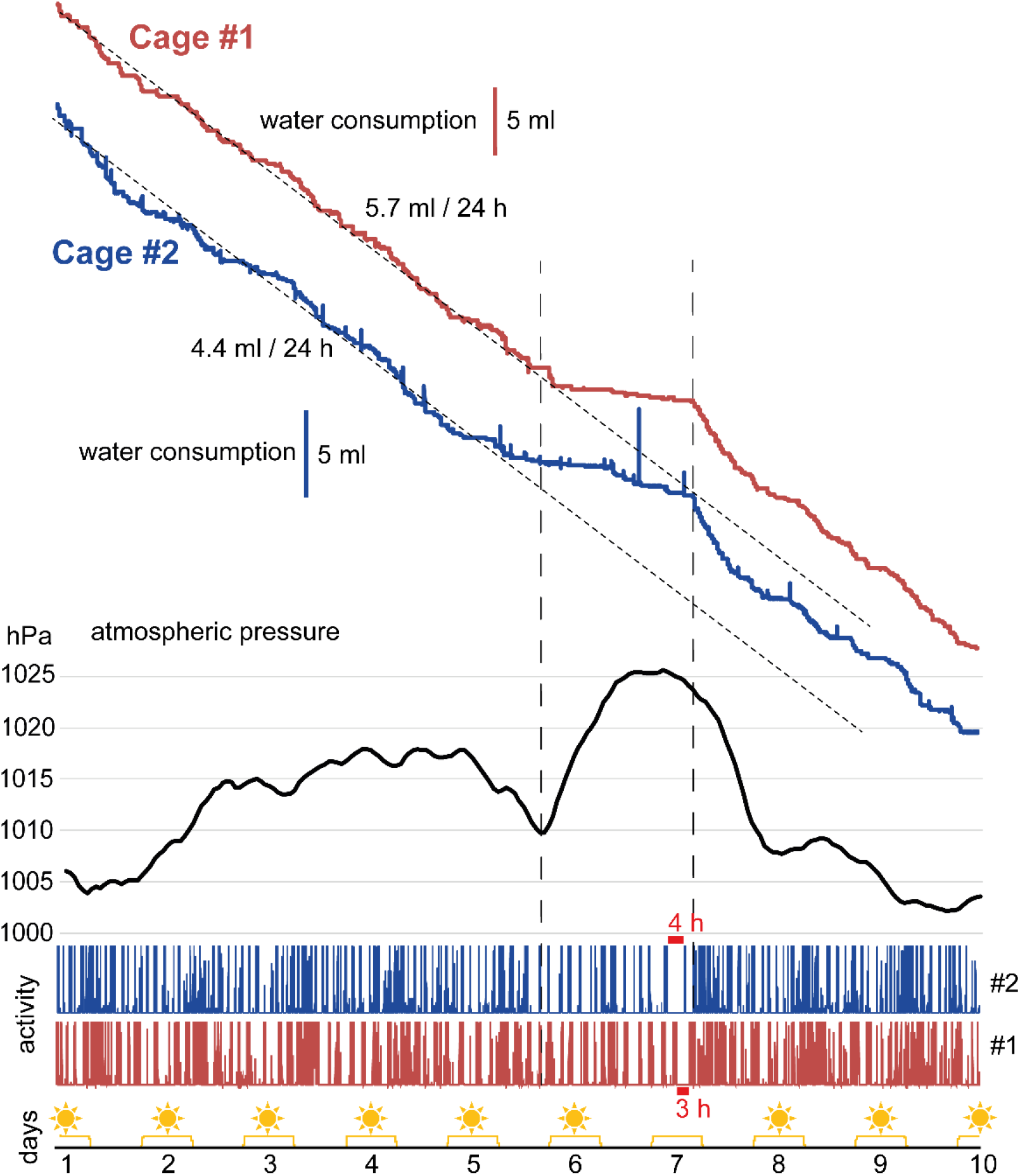
Reduced water consumption in two mouse cages during period of increasing atmospheric pressure. Water consumption in two cages (1 mouse each, standard water bottle). On the second day of reduced water consumption (day 7), mouse #1 is motionless for 3 h and mouse #2 is motionless for 4 h (red bars). Light periods indicated in yellow.

To investigate the interplay between air pressure and bubble formation more closely, we replaced the cylindrical section of the sipper tube with a borosilicate glass tube of the same dimensions, keeping the original tip with the drinking hole. We connected a modified syringe pump to the air-filled space at the top of the water bottle via a flexible PVC tube (Fig. 3A). To simulate rising atmospheric pressure, we slowly reduced the pressure inside the bottle (10 ml/h). At a pressure differential of about 3 hPa to the outside air, the surface tension at the drinking hole was overcome, resulting in the periodic formation of air bubbles during constant suction (Fig. 3B). Each air bubble formed rapidly, then slowly moved upwards inside the sipper tube (Fig. 3C). These “Taylor bubbles” are a well-known fluid dynamics phenomenon; if they become trapped, an air lock is created^4^. We reasoned that a coiled wire inside the sipper tube would create a path for water to flow under the bubble and dislodge it. Indeed, this modification increased the speed that bubbles move up the tube threefold (Fig. 3D, E). The size of the bubbles, measured as the differential pressure relieved per bubble (step size in panel B), was not affected by the insert (Fig. 3F).

**Fig. 3:**
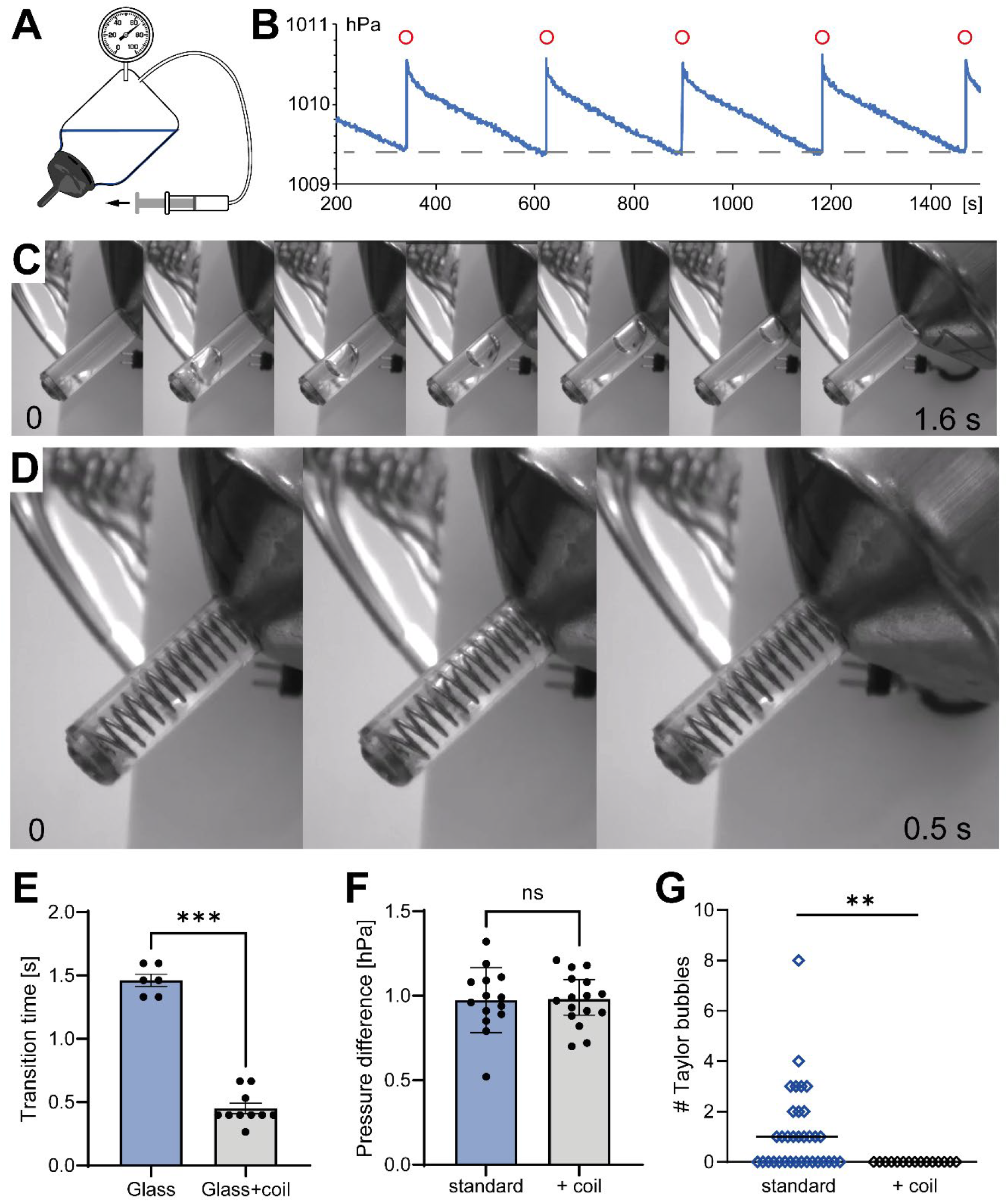
Wire coil insert prevents air locks. **A)** Experimental apparatus to monitor bottle pressure during continuous suction via syringe pump. **B**) Whenever the bottle pressure drops below threshold (dashed line), an air bubble is generated and rises in the bottle (red circles). Each bubble reduces the pressure differential by ∼1 hPa, generating a sawtooth pattern. Atmospheric pressure was 1012 hPa. **C**) Movie frames of a spontaneous pressure equalization event. A Taylor bubble forms rapidly and moves slowly through the sipper tube. **D)** The same glass tube with wire coil insert. The bubble rapidly passes through the coil in a single movie frame (7.5 fps). **E)** Coil insert significantly speeds up bubble transition. Bars show mean ± SEM. n = 6, 10 events. Welch’s t test, ^***^p < 0.0001. **F**) Bottle pressure drop (before/after bubble event) is not affected by coil insert. Bars show mean ± SEM. n = 14, 16 events. Welch’s t test, p = 0.99. **G**) Coil insert reduces the probability of trapped bubbles in metal sipper tubes to zero. Each open diamond corresponds to a bottle (n = 34, 15 bottles), 30 daily measurements each. Horizontal lines show median. Kolmogorov-Smirnov test, p = 0.0059.

Next, we tested an array of 49 bottles daily for trapped bubbles. Bottles were equipped with standard sipper tubes (AISI 316 stainless steel), some of which contained coil inserts that were not visible from the outside. Over an observation period of 30 days, daily blind measurements revealed trapped bubbles (on one or more days) in more than half (18/34) of the standard sipper tubes, while tubes with coil inserts never contained air bubbles (Fig. 3E). In two of bottles with standard sippers, we observed cases where the meniscus had completely retracted, leaving no water in the first 20 mm behind the drinking hole. This did not happen in bottles with a coil insert. These experiments revealed that air locks and meniscus retraction are not rare phenomena in standard metal bottle caps, but are completely prevented by a wire coil insert.

Mice are very sensitive to dehydration^2,5^. Acute water deprivation for 24 h leads to 18-20% loss of body mass^6^. When access to water is restricted to 15 minutes per day, physical growth and cognitive development of young mice is severely impaired^7^. Based on our measurements, we believe that weather-induced dehydration (Fig. 2) may affect a large number of laboratory animals. While fatalities are rare, dehydration introduces a significant, uncontrolled confounding factor. Unknown to the experimenters, some animals may enter experiments dehydrated, which is known to alter their behavior and metabolism, e.g. responses to drugs^5^.

As mouse cages and drinking bottles are internationally standardized, we assume that more than one facility or country is affected. The sporadic malfunctioning of water bottles is a known problem that has been blamed on carbonate deposits, greasy residue, or other forms of contamination^3,8^. A casual inspection of suspect bottle caps inevitably focuses on the small tip opening (1.8 mm diameter) and the possibility of clogging. This line of reasoning permeates the guidelines of the Society of Laboratory Animal Science^3^. We suggest that air locks in drinking tubes are a heretofore unappreciated factor contributing to water bottle failure. The physics of Taylor bubbles involves a delicate balance between buoyancy, surface tension, and viscous forces, which together create a stable, non-moving state^9,10^. The surface of stainless-steel forms a hydrophobic oxide layer (Cr_2_O_3_) that promotes Taylor bubble stability. Coating the inside of the sipper tube with hydrophilic polymers or TiO_2_ could alleviate the problem^11,12^. However, animal facilities use hot water combined with acidic or alkaline cleaning agents to clean drinking bottles and caps, followed by sterilization in an autoclave. The resistance of thin coatings to harsh cleaning cycles is unclear. Strongly conical bottle caps (e.g. Allentown 223402) also prevent airlocks, but are not compatible with all cage types.

We show that a coiled wire insert disrupts the boundary layer between the bubble and the tube, creating a passageway for downward flow which dislodges the bubble^4^. When integrated during production, the coil insert has an additional advantage in that it allows for the use of a fully open sipper tube (5.5 mm tip opening) without leaking, thus reducing manufacturing costs while minimizing sipping effort.

## Methods

### Animals

Adult C57BL/6 mice of either sex were bred in the animal facility of the University Medical Center Hamburg-Eppendorf. We received permission for the animal experiments from the local authorities (Behörde für Justiz und Verbraucherschutz der Freien und Hansestadt Hamburg, Permit N110/2020). All procedures were in accordance with EU guidelines for animal experiments (Directive 2010/63/EU).

### Laboratory animal equipment

We used standard mouse cages (Tecniplast 1284), polycarbonate water bottles (400 ml, Tecniplast ACBT0402) and standard bottle caps (AISI 316 stainless steel, 1.8 mm hole, Tecniplast ACCP2521) taken from the circulation at the animal facility of the University Medical Center Hamburg-Eppendorf. All tested water bottles and bottle caps had been in use for years; we therefore make no claims about factory-new products.

Wire coils were manufactured in-house from 0.6 mm diameter stainless steel wire with an outer diameter of 5.1 mm and a length of 25 mm. Coils and modified bottle caps are commercially available from corresponding author F.K. on request.

### Testing apparatus

To detect bubble transition, we equipped standard mouse bottles with a pressure sensor (MS5837 02BA, TE Connectivity) inside the metal cap (Fig. 1). Pressure was sampled at 1 Hz using a custom ATxmega256A3 electronic board. Every minute, the min-max difference was calculated over the last 60 data points and saved. We visually confirmed that the pressure spikes correspond to air bubbles passing the sensor inside the cap. Tongue contacts, indicative of licking events, were detected by monitoring the resistance between the metal cage floor and the bottle cap. Every second with lick events was scored as ‘active’; the sum of active seconds per minute is plotted in Fig. 1. The same sampling strategy was used to score proximity events with a pyroelectric sensor (Napion AMN33111J, Panasonic). Data were graphed with MS Excel 2019.

To measure water consumption over long time periods with a precision of ±20 ml (Fig. 2), we used Nutrition Monitor cages (INFRA-E-MOTION) with a custom 45° bottle holder and passive IR motion sensor (Napion AMN33111J, Panasonic). Air pressure data for Hamburg (Finkenwerder West) were downloaded from https://hamburg.luftmessnetz.de/meteorology/ldr.

To detect bubble formation (pressure equalization events, Fig. 3), we measured air pressure inside the bottle at the highest point of the 45° downward tilted bottle (MS5837 02BA, TE Connectivity). Constant suction was provided by a modified syringe pump (Al-1000, Aladdin). For optical measurements of Taylor bubble speed, transparent sipper tubes were made from borosilicate glass sections sealed between the tip and the root of a steel sipper tube with cyanoacrylate glue. Movies were recorded with a 5 MP industrial CMOS camera (The Imaging Source). To detect air bubbles in steel sipper tubes, we slowly advanced a thin enameled copper wire through the drinking hole in axial direction (45°) while measuring the resistance to the bottle cap. 49 bottles were arranged in 7 racks, coils were inserted before the start of the test in a random subset of bottle caps. A sudden drop in resistance, indicated by a light signal (LED), indicated that the tip of the wire had entered an air pocket.

We used GraphPad Prism 10 for bar plots and statistical analysis (Welch’s t test, two-tailed; Kolmogorov-Smirnov test).

## Acknowledgements

The authors would like to thank the late Katja Husen for continuous support of the project. We thank the animal care technicians and animal welfare officers that informed us about the dehydration problems, but wish to remain anonymous.

## Contributions

F.K. designed the study, collected and analyzed the data and revised the manuscript. S.L. supported data collection and revised the manuscript. T.G.O drafted the manuscript. All authors edited the final version of the manuscript.

## Supplemental Material

**Supplemental Figure 1:**
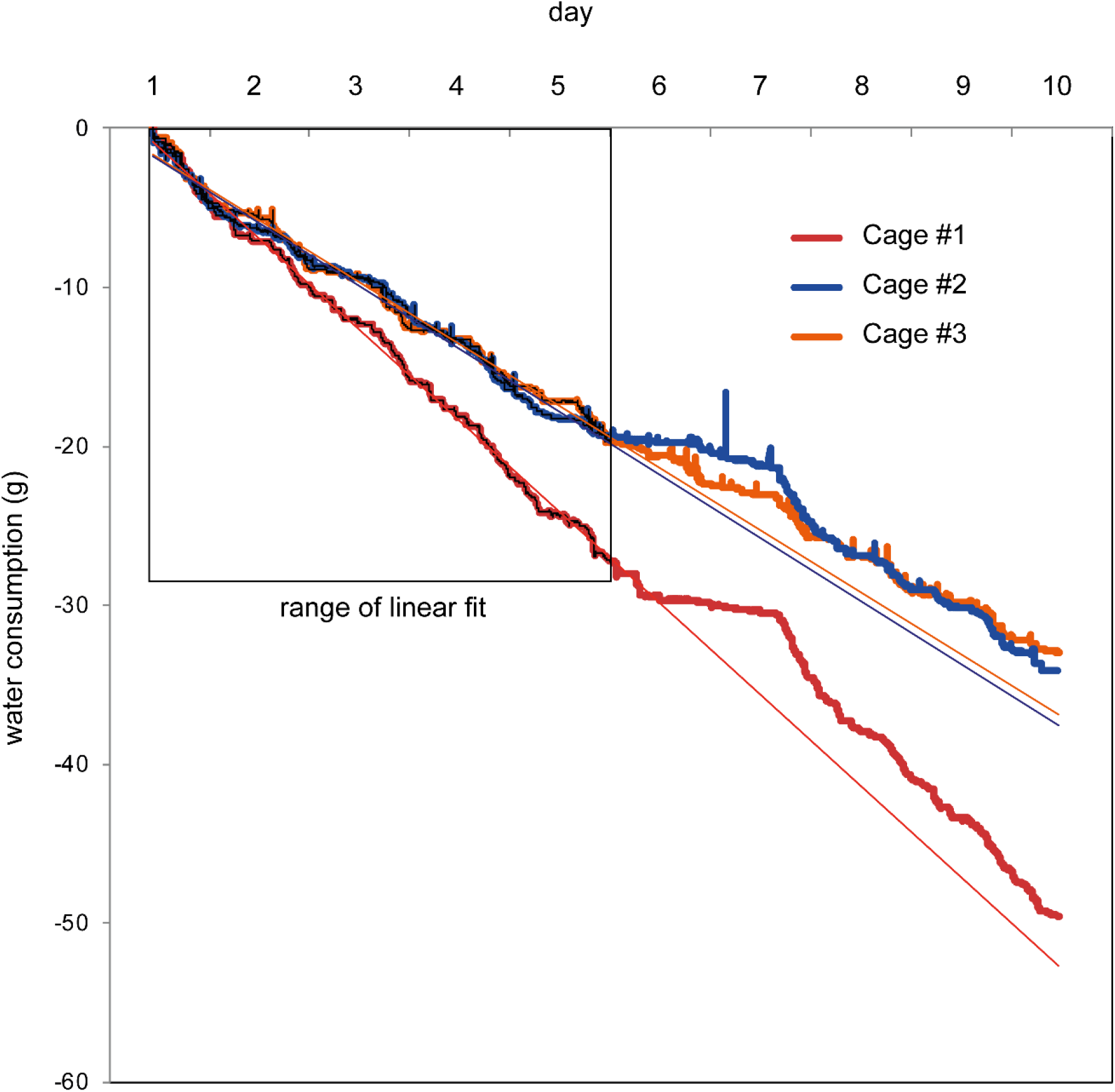
Linear extrapolation of water consumption on the first 5 days reveals synchronized periods of reduced consumption (day 6 and 7) in three out of nine cages, followed by enhanced drinking over several hours. Data from cage #1 and #2 data are identical to Fig. 2.

## References

1. Animal Testing: Decline of Previous Years Clearly Continues. https://www.bfr.bund.de/en/press-release/animal-testing-decline-of-previous-years-clearly-continues/ (2024).

2. Bekkevold, C. M., Robertson, K. L., Reinhard, M. K., Battles, A. H. & Rowland, N. E. Dehydration parameters and standards for laboratory mice. J. Am. Assoc. Lab. Anim. Sci. 52, 233–239 (2013).

3. I. Hagelschuer H. Wagner G. R. Warncke. Trinkwasserversorgung von Versuchstieren. https://www.gv-solas.de/wp-content/uploads/2021/08/2016Trinkwasserversorgung.pdf (2016).

4. Zhou, G. & Prosperetti, A. Faster Taylor bubbles. J. Fluid Mech. 920, (2021).

5. Rowland, N. E. Food or fluid restriction in common laboratory animals: balancing welfare considerations with scientific inquiry. Comp. Med. 57, 149–160 (2007).

6. Schwartz, N. E., Alva, M. R. & Garland, T., Jr. Differential effects of acute total water deprivation on voluntary exercise behavior and body mass in laboratory house mice. Physiol. Behav. 303, 115139 (2026).

7. Kim, C.-S., Chun, W. Y. & Shin, D.-M. Dehydration impairs physical growth and cognitive development in young mice. Nutrients 12, 670 (2020).

8. Working Group on Cage Processing. Cage Processing in Animal Facilities. https://felasa.eu/Portals/0/7_%20Auflage%20AK%20KAB-Broschure_2024_e.pdf (2024).

9. Balestra, G., Zhu, L. & Gallaire, F. Viscous Taylor droplets in axisymmetric and planar tubes: from Bretherton’s theory to empirical models. Microfluid. Nanofluidics 22, (2018).

10. Dhaouadi, W. & Kolinski, J. M. Bretherton’s buoyant bubble. Phys. Rev. Fluids 4, (2019).

11. Gatou, M.-A., Syrrakou, A., Lagopati, N. & Pavlatou, E. A. Photocatalytic TiO2-based nanostructures as a promising material for diverse environmental applications: A review. Reactions 5, 135–194 (2024).

12. Garcia-Perez, V. I. et al. Amorphous TiO2nano-coating on stainless steel to improve its biological response. Biomed. Mater. 19, 055037 (2024).

